# Mutualistic acacia-ants show that specialized bacteria are not required for the evolution of herbivory

**DOI:** 10.1101/208215

**Authors:** Benjamin E.R. Rubin, Stefanie Kautz, Brian D. Wray, Corrie S. Moreau

## Abstract

Acacia-ant mutualists in the genus *Pseudomyrmex* nest obligately in acacia plants and, through stable isotope analysis, we show that they are among the strictest of herbivores, feeding exclusively from their hosts. The diets of herbivorous insects such as these are often enriched by obligate bacterial endosymbionts through nitrogen recycling and even gaseous di-nitrogen fixation. We, therefore, examine the bacterial communities associated with mutualistic acacia-ants, comparing them with related non-mutualists in order to determine whether they host bacterial partners likely to contribute to the enrichment of their diets. However, despite their low trophic position, we find no evidence for bacteria-assisted nutrition in either adults or larvae. These acacia-ants do not host any species- or clade-specific bacteria, though several lineages of acetic acid bacteria present across social insects do differ in abundance between mutualists and non-mutualists, likely in response to the sugar-rich diets of their hosts. In addition, two novel lineages of Actinomycetales inhabit both mutualistic and non-mutualistic *Pseudomyrmex* and differ in abundance between the juveniles of these groups, potentially serving as defensive symbionts. Metagenomic sequencing of these taxa reveal substantial capacity for the production of defensive chemicals. Overall, we find little evidence for nutrition-associated bacteria in these strictly herbivorous ants, showing that bacteria are not as essential to animal nutrition as is often hypothesized.

## Introduction

Animals often depend on endosymbiotic bacteria to digest, process, and enrich their diets (Flint, Bayer, Rincon, Lamed, & White, 2008; Hansen & Moran, 2011; Moran & Baumann, 2000). Some of the best understood cases of animal dependence on bacteria for dietary enrichment are from insects. Termites, for example, depend on endosymbiotic bacteria to fix nitrogen (Benemann, 1973; Breznak, Winston, Mertins, & Coppel, 1973) and pea aphids (*Acyrthosiphon pisum*) depend on the obligate intracellular symbiont *Buchnera aphidicola* to produce amino acids without the need for dietary input beyond nitrogen-poor plant phloem (Hansen & Moran, 2011; International Aphid Genomics Consortium, 2010; J. P. Sandström & Moran, 2001; J. Sandström & Pettersson, 1994). The strict cospeciation between aphids and their *Buchnera* endosymbionts as well as the maternal deposition of bacteria in embryos further support the tight relationship between these organisms (Miura et al., 2003). Coevolution between a variety of other insects and their endosymbionts (e.g. Bandi et al., 1995; Cheng & Aksoy, 1999) show that these relationships are not unusual.

Recently, ants have become a target of research into the relationships between insects and bacteria (Funaro et al., 2011; Hu et al., 2017; Hu, Łukasik, Moreau, & Russell, 2014; Łukasik et al., 2017; Moreau & Rubin, 2017; Ramalho, Bueno, & Moreau, 2017; Russell et al., 2009; Sanders et al., 2014). These insects were long thought to be predominantly predatory (Floren, Biun, & Linsenmair, 2002) but this hypothesis has been challenged on the basis that these animals drastically outnumber other arthropods in a variety of ecosystems (Davidson, Cook, Snelling, & Chua, 2003; Davidson & Patrell-Kim, 1996; Majer, 1990). Specialization on nitrogen poor plant exudates has emerged as the most likely scenario (Davidson et al., 2003). However, these carbohydrate rich but protein poor resources appear not to provide the amino acids required for colony growth and reproduction (Davidson, 1997), leaving the question of ant nutrition unanswered, though dependence on gut bacterial enrichment of diet through nitrogen enrichment is a compelling possibility (Davidson et al., 2003; Feldhaar et al., 2007; Russell et al., 2009; Zientz, Feldhaar, Stoll, & Gross, 2005). The ant genus *Cephalotes*, a strict herbivore, has provided some of the first suggestions that this is the case, hosting codiversified bacterial communities rich in Rhizobiales, the same bacterial order responsible for fixing gaseous nitrogen into a form usable by legumes (Hu et al., 2014; Russell et al., 2009; Sanders et al., 2014). Relationships with Rhizobiales have also been identified in *Tetraponera*, suggesting that this symbiosis may be widespread in ants (Borm, Buschinger, Boomsma, & Billen, 2002; Stoll, Gadau, Gross, & Feldhaar, 2007). While these findings are compelling, no functional assays of Rhizobiales in these species have been completed.

The obligate mutualism between ants in the Neotropical genus *Pseudomyrmex* and acacia trees is one of the best known of any ant-plant relationship (Janzen, 1966, 1967). In this symbiosis, ants nest in and feed on the hollow thorns, food bodies, and extra-floral nectar provided by the plant. In exchange for these resources, resident ants aggressively protect their hosts by attacking herbivores, trimming encroaching plants, and removing pathogenic fungi (Janzen 1973). While many herbivorous ants are opportunistic predators and animal-protein scavengers, *Pseudomyrmex* mutualists appear to have an unusually strict plant-based diet. Clement, Köppen, Brand, & Heil (2008) presented animal-protein in a variety of forms to nests of the mutualist *P. ferrugineus* and, rather than consume these protein-rich resources, the ants actively rejected them, dropping them from their host trees. *Pseudomyrmex gracilis*, a sympatric non-mutualistic congener, instead collected and, presumably, ate the available protein sources. The range of resources utilized by non-mutualistic *Pseudomyrmex* species is clear from stable isotope studies that classify them as both herbivores and predators, depending on environment (Tillberg, Holway, LeBrun, & Suarez, 2007). The strictly herbivorous acacia-ant diet coupled with the omnivorous diets of their congeners make this group ideal for studying the evolution of ant herbivory. Indeed, previous work has suggested that associations with nitrogen-fixing bacteria may exist (Eilmus & Heil, 2009).

In order to determine whether bacteria are likely contributing to the nutrition of mutualistic *Pseudomyrmex* acacia-ants, we used 16S rRNA amplicon sequencing to characterize the microbial communities within these species. We also determined the specificity of these relationships by examining sympatric non-mutualistic *Pseudomyrmex* as well as ants from 11 other distantly related genera. Finally, we confirm the degree to which acacia-ants are strictly herbivorous, as has been previously proposed, by estimating ratios of nitrogen isotopes. Together, these data show the connection between plant-ant mutualism, trophic ecology, and bacterial community composition.

## Materials and Methods

### Sampling

All ant samples were collected from the Area de Conservación Guanacaste (ACG) in northwestern Costa Rica at the Santa Rosa Biological Station (10.8° N, 85.6° W) during June of 2012 and from the Florida Keys, USA (25.1° N, 80.5° W) between 2011 and 2014. Within the ACG, there were four collecting sites that were used: Tanqueta trail, Barrachas trail, River trail, and Playa Naranjo. Three species of acacia-ant mutualists were collected from the ACG: *Pseudomyrmex flavicornis*, *P. nigrocinctus*, and *P. spinicola*. Each of these species nests obligately in acacia (genus *Vachellia*) plants and behave as mutualistic partners. *Pseudomyrmex gracilis*, a widely distributed non-mutualist (generalist) co-occurs in this area and was also a focal taxon for collection to allow for comparisons between mutualists and closely related generalists. Although rare, we also collected *P. nigropilosus* whenever possible. This species is more closely related to *P. gracilis* than to acacia-ants but nests obligately in acacias, acting as a parasite on the mutualism, providing no protection to the host plant (Janzen, 1975). Plant-ant mutualism has convergently evolved three times within *Pseudomyrmex* (Chomicki, Ward, & Renner, 2015; Rubin & Moreau, 2016; Ward & Downie, 2005), with three types of plants: acacias, *Triplaris*, and *Tachigali*. A single Triplaris-nesting species, *P. viduus*, also exists in the ACG and was collected when possible. Finally, we also included a single unidentified *Pseudomyrmex* generalist found incidentally.

No mutualistic species range into the Florida Keys but several *Pseudomyrmex* generalists do occur there, including *P. gracilis*. This species was again a focal point for collection so as to allow within species comparisons across a large geographic distance. In addition, *P. cubaensis, P. elongatus, P. simplex, P. seminole*, and *P. pallidus*, all generalists, were also collected from this site.

In both areas, we also performed general collections of ants, focusing in particular on the three taxa that are known to host bacterial symbionts (*Cephalotes, Camponotus*, and Dorylinae, the army ants) though all taxa were included. We found that many acacias were also parasitized by a species of *Crematogaster* and we collected this species wherever possible. In general, we made collections from at least three colonies of each target taxon from each of the local collecting sites in Costa Rica.

Lastly, we also included collections of known herbivores (e.g. caterpillars, termites, and millipedes), carnivores (e.g. spiders and centipedes), and plants, including a large number of acacias, so as to calibrate the stable isotope analyses. All collections were made into 95% alcohol in the field and kept at −20° C until processing. All samples used in this study are shown in Table S1 and vouchers have been deposited in the scientific collections of the Field Museum of Natural History, Chicago, Illinois, USA.

### Trophic levels

We collected a total of 359 samples for stable isotope analysis. These included leaves from 59 acacia plants occupied by *Pseudomyrmex* mutualists, ants from 54 acacia-ant mutualist colonies, 12 colonies of the generalist *P. gracilis* from the same sites in Costa Rica as the mutualists, 40 colonies of *Cephalotes*, as well as 30 known herbivores and 15 predators. These samples were collected from all four ACG sites. Additional generalist *Pseudomyrmex* and co-distributed insects from Florida were also included.

We pooled between three and 20 heads and thoraces (mesosomas) of adult ants from individual colonies for each isotope measurement. Samples were dried before analyzing. For one colony of *P. nigropilosus*, the acacia parasite, we did not have sufficient adults and instead used pupae. Plants were collected in the same fashion and ~1cm^2^ of leaf material was used to estimate isotope ratios. All samples were analyzed at the University of Utah Dartmouth Stable Isotope Ratio Facility for Environmental Research (Salt Lake City, Utah, USA). For other insects, segments containing guts were removed when possible. For caterpillars, millipedes, and centipedes, the front half of the insect was typically used.

For each site in the ACG, we had at least 11 samples of acacia leaves. We, therefore, standardized all ACG samples to the mean δ^15^N ratio of the acacias at the same site. We then used FDR-corrected Wilcoxon rank-sum tests to compare nitrogen ratios between taxa. To make the samples from the Florida Keys comparable to those from Costa Rica, we subtracted the difference in median values measured for all plants sampled in Costa Rica from the median value for plants from the Florida Keys. We used the median because the spread of plant values was quite large. We, again, used Wilcoxon rank-sum tests to compare between taxa.

### DNA extraction

All DNA extractions were performed using the Mo Bio PowerSoil Kit with minor modifications (Rubin et al., 2014). Abdomens (gasters) were removed and surface sterilized in 5% bleach for one minute before DNA extraction. Gasters were used in an attempt to enrich for gut-associated microbes. For the larger species, including all *Pseudomyrmex*, a single gaster was included in each extraction, though multiple gasters were pooled for the smaller species. Several larvae were pooled whenever possible. Four blank samples with no insect material were also included in DNA extraction and subsequent sequencing to aid in the identification of contaminant sequences.

### Bacterial quantification

Quantification of the bacterial 16S rRNA gene was done as previously (Rubin et al., 2014). We required that all qPCR reactions had efficiencies and R^2^ between 90% and 110%. Each sample was run at least three times and, when one of these was eliminated for quality control, run a fourth time. The mean was taken when multiple values were available. QPCR results were standardized to the overall quantity of DNA measured using a Qubit fluorometer (Rubin et al., 2014). Differences in quantity of bacteria between taxa was assessed using Wilcoxon rank-sum tests with FDR-correction for multiple testing.

### Bacterial 16S rDNA sequencing

Sequencing was done following the protocol of the Earth Microbiome Project (Caporaso et al., 2012). The bacterial 16S rRNA gene was amplified using the universal primers 515f (5′ - GTGCCAGCMGCCGCGGTAA) and 806r (5′ - GGACTACHVGGGTWTCTAAT) and sequenced using two lanes of Illumina MiSeq 150bp paired-end sequencing.

Despite the PCR amplification inherent in amplicon sequencing of the bacterial 16S rRNA gene, successfully obtaining informative data from this procedure is challenging, particularly when the concentration of bacterial DNA is low, as is known from many ants (Hu et al., 2017; Rubin et al., 2014; Sanders et al., 2017). In order to guarantee that we would obtain data from a sufficient number of colonies, we, therefore, performed three DNA extractions from each colony for both adults and larvae whenever possible. Overall, we sequenced adults and larvae from 176 ant colonies from 11 ant genera, resulting in 589 sequencing samples.

### 16S rDNA sequence processing

The two lanes of sequencing results were pooled into a single dataset for all analyses. 16S rRNA gene sequences were demultiplexed using QIIME (Caporaso, Kuczynski, et al., 2010; Navas-Molina et al., 2013) and then forward and reverse reads were merged using UPARSE (Edgar, 2013), truncating at the first base with quality of three or less. The resulting contigs were then filtered for quality by discarding reads with more than 0.5 expected errors using the fastq_filter function of UPARSE.

Contamination can be a major issue for microbiome studies based on bacterial 16S sequencing. We therefore used Sanders’s decontamination pipeline to remove probable contaminants from our dataset (https://github.com/tanaes/decontaminate). We first clustered OTUs within our four blank samples at 97% similarity and then discarded all OTUs that made up less than 0.1% of these blank sample communities. The most abundant sequences in each of these OTUs were then used as seeds to identify all other sequences of 97% similarity which were deemed to be contaminants and discarded. We then used the default UPARSE pipeline to cluster sequences into OTUs, discarding those represented by fewer than two sequences and filtering chimeras with the gold database provided by UCHIME. Sequences were clustered at 97% similarity. Fasttree as implemented in QIIME was used to infer a phylogeny for all representative sequences.

We filtered all samples with fewer than 1,000 sequences after quality controls. We split all ant samples into two sets of adults and larvae. In many cases, we sequenced multiple individuals from each colony. We combined these non-independent samples into single representatives of colonies by summing reads in all individuals from each colony.

We tested for differences in bacterial communities using several approaches. First, we used supervised learning as implemented in QIIME to determine if communities are identifiable as originating from particular types of ants. We ran these analyses on 100 replicate OTU tables rarefied to 1,000 reads and used the average error ratio to assess their performance. We used an error ratio (the baseline error rate if the classifier worked randomly to the observed errors) ≥ 2 to determine when significant differences in communities exist (Navas-Molina et al., 2013; Van Treuren et al., 2015). We also compared beta diversities (measured as weighted and unweighted Jaccard and UniFrac distances) within and between groups to determine how consistent communities were, testing for differences using non-parametric t-tests with Monte Carlo permutation corrected for multiple testing using the Benjamini-Hochberg procedure. Lastly, we also tested for differences in the abundance of individual taxa using Wilcoxon rank-sum tests and the frequency of the presence of individual taxa using *χ*^2^ tests both corrected for multiple testing using the Benjamini-Hochberg procedure.

### Ant specificity

It is clear that *Pseudomyrmex* ants are occupied by a wide variety of bacterial taxa, including many widespread bacteria picked up from the environment and contaminants from human handling. We first attempted to determine whether taxa were present in the wider environment by also sequencing bacteria drawn from sympatric plants. However, many potentially interesting taxa were present at vanishingly low levels on these plants - on the order of one or two sequences. This number of reads could very easily result from laboratory contamination between samples but was confounding when attempting to identify environmental contaminants.

Therefore, we produced a further reduced set of target OTUs by clustering the Earth Microbiome Project's 2013 10k representative sequence dataset against our *de novo* OTUs. This dataset includes sequences from more than 13,000 samples from a variety of environments and materials, including Neotropical forests, and should include representatives of most widespread bacterial taxa. Those of our core OTUs that did not cluster at 97% with any EMP sequences were assumed to be novel taxa and likely ant or insect specific. There were 805 of these OTUs out of an initial 5,356 making up 2,804,128 sequences.

For every set of OTUs of interest (e.g. those significantly different in abundance between adults and larvae), we tested whether these ant-specific OTUs were overrepresented using hypergeometric probabilities of observing the given number of ant-specific OTUs in the current set of interest. Each set was compared to the background of 805 ant-specific OTUs out of a total of 5,356.

In order to determine just how unique our set of dominant ant-specific OTUs were, we inferred the phylogenetic history of those ant-specific OTUs that were present in at least 10 samples. For each OTU, we also included the closest BLAST hit in NCBI's nucleotide database. The phylogeny was inferred using RAxML (Stamatakis, 2006) with the GTRGAMMA model of nucleotide substitution.

### Metagenomes

To assess the functional capacity of the bacterial communities associated with *Pseudomyrmex*, we performed shotgun metagenomic sequencing of two libraries constructed with Illumina's Nextera XT kit. The first sequencing library, which included DNA extracted from 25 larval meconia from *P. nigropilosus* colony BER0554, was sequenced using 150bp paired-end reads on the MiSeq platform. The second library, with DNA from 30 adult guts from *P. flavicornis* colony BER0517, was sequenced using two lanes of HiSeq paired-end 100bp sequencing. We chose these colonies based on available material and the relative abundances of bacteria of interest.

The resulting sequences were filtered extensively. We first removed duplicate reads using FastUniq (Xu et al., 2012). The remaining sequences were filtered using Trimmomatic (Bolger, Lohse, & Usadel, 2014) with the following parameters: LEADING:3 TRAILING:3 SLIDINGWINDOW:4:10 MINLEN:36. In addition, given the frequent contamination of Nextera libraries with Illumina adapter sequences, we aggressively filtered adapters with ILLUMINACLIP:2:25:7. This filtering introduced a small number of additional duplicate sequences (by trimming errors) so we reran FastUniq on the filtered sequences. We then merged paired reads into single reads using fastq-join from the ea-utils package.

The resulting unpaired sequences as well as the remaining paired-end sequences were mapped to the *P. gracilis* genome (Rubin & Moreau, 2016) using Stampy v. 1.0.23 (Lunter & Goodson, 2011) allowing 3% divergence between reads and reference. Reads that failed to map were used in all subsequent analyses. If one read in a pair was successfully mapped, both reads in that pair were discarded. We performed analyses using the MG-RAST webserver.

## Results

### Acacia-ants have an extremely low trophic level

Acacia-ants have a significantly lower trophic level than every other ant genus sampled as well as the non-mutualistic *Pseudomyrmex* species *P. gracilis* and the *Triplaris*-inhabiting mutualist *P. viduus* (FDR-corrected P < 0.05, Table S2, Fig. 1). The only taxa for which acacia-ants do not have significantly lower δ^15^N ratios are the acacia-parasite *Pseudomyrmex* species, *P. nigropilosus*, and a generalist collected in the Florida Keys, *P. cubaensis* (Fig. 1).

**Figure 1.**
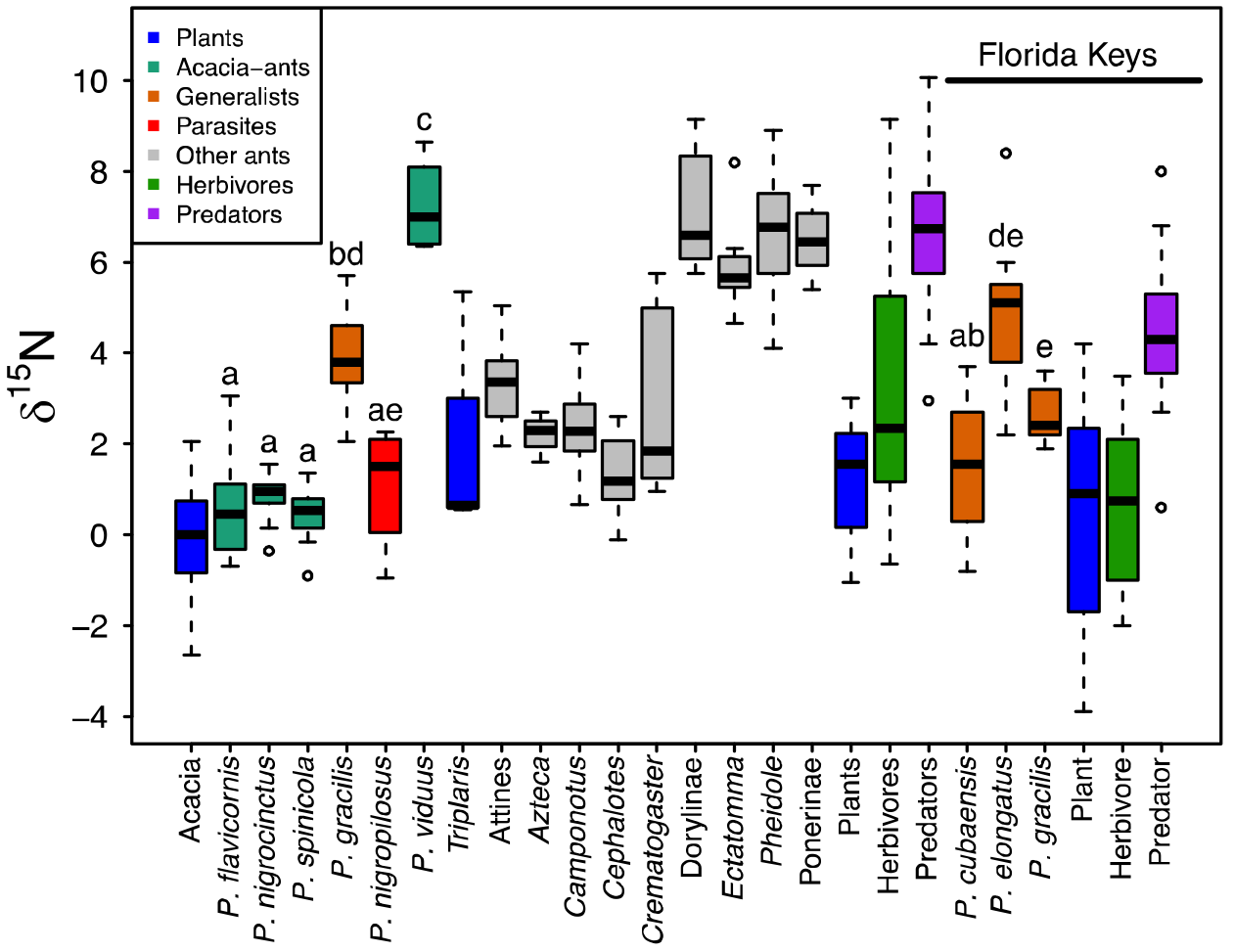
Distribution of δ^15^N across samples collected in the ACG and the Florida Keys. Letters show significance between ant species of *Pseudomyrmex* based on Wilcoxon rank-sum tests with FDR-correction. All three acacia-ant species (P. *flavicornis, P. nigrocinctus*, and *P. spinicola*) also have significantly lower δ^15^N than every ant taxon included in this study. *Pseudomyrmex gracilis* is a generalist, *P. viduus* is a plant-ant mutualist that nests obligately in trees in the genus *Triplaris*, and *P. nigropilosus* is an obligate parasite of acacia plants.

### Bacterial quantity varies by genus

The quantity of bacteria varies widely in adults by ant genus (Fig. 2). Notably, those taxa with clearly established symbiotic (and likely mutualistic) relationships with endosymbiotic bacteria (*Camponotus* and *Cephalotes*) have the highest number of bacteria and significantly more than all species of *Pseudomyrmex*, regardless of collection site (P < 0.0006).

**Figure 2.**
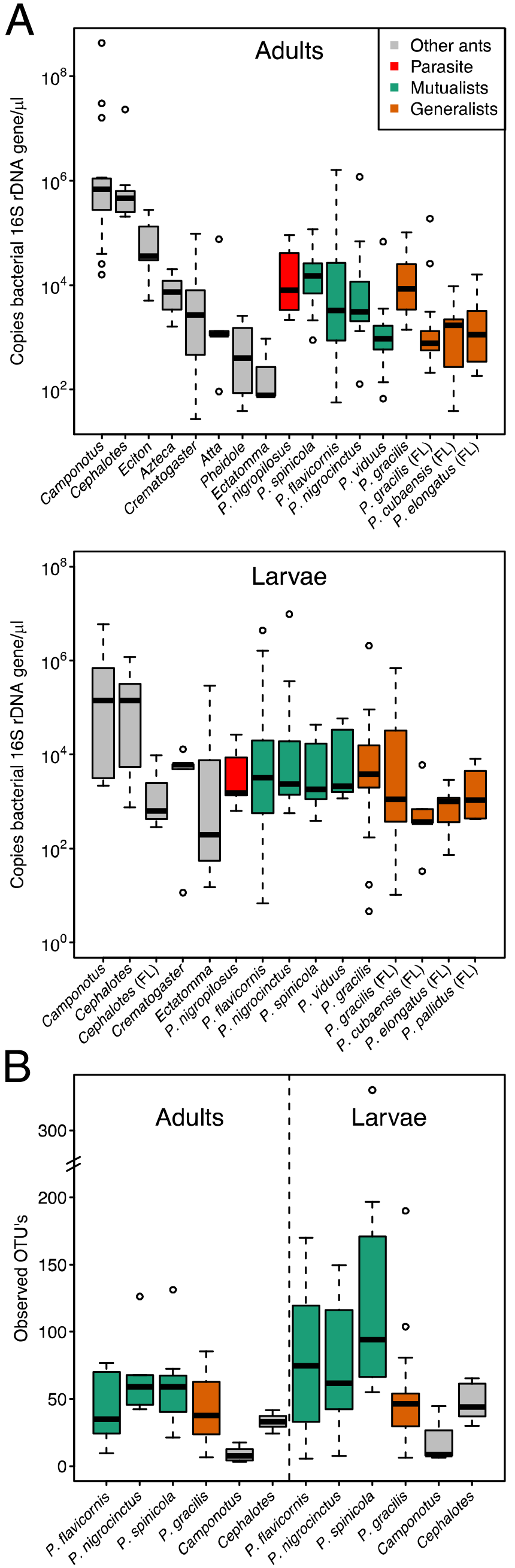
Quantities of the bacterial 16S rRNA gene in the ant taxa examined using qPCR in adults and larvae (A) and alpha diversity in specific species and genera collected in the ACG (B). All *Pseudomyrmex* adults, including obligate mutualists, have significantly lower numbers of bacteria than both *Camponotus* and *Cephalotes*, the two genera with known beneficial bacterial symbionts. Significance was determined by Wilcoxon rank-sum tests with FDR-correction (P < 0.05).

There is little variation in quantity of bacteria present between species of *Pseudomyrmex* collected from the same site (Fig. 2A, Table S3). Between the adults of the five species collected from the ACG, *P. viduus* has significantly less bacterial DNA than all other taxa (P < 0.05) except *P. flavicornis* (P = 0.2). No other significant differences were identified (P > 0.05). However, excluding *P. viduus*, all *Pseudomyrmex* species collected from the ACG had significantly greater numbers of bacteria than all species collected from Florida (P < 0.05). Notably, this includes the difference in quantity between *P. gracilis* collected from each site (P = 0.0003), strongly indicating that locality impacts quantity of bacteria present, at least over the large distances studied here. *Pseudomyrmex viduus* is not significantly different to any of the taxa from Florida (P > 0.05).

There is far less variation within larvae, though the same general patterns do hold (Fig. 2). *Camponotus* has significantly greater numbers of bacteria than all *Pseudomyrmex* taxa except *P. pallidus*, though these differences do not exist for *Cephalotes*. The significantly lower numbers of bacteria in *Pseudomyrmex* from Florida compared with those from the ACG, while not as consistent as for adults, occur for the majority of comparisons, including for the comparison of *P. gracilis* collected at the two sites (P < 0.05).

### 16S sequencing

We obtained a total of 27,035,218 16S rRNA sequencing reads. Of these, 4,994,866 could not be assigned to samples due to sequence errors in barcodes. Quality filtering left 17,877,523 and contaminant filtering left 11,151,725 high quality, contaminant filtered sequences. There were 1,290,634 unique sequences and we discarded all singletons from this set, leaving 304,342 seed sequences. 29,732 of these were identified as chimeras by the clustering step of UPARSE and the remaining sequences clustered into 8,131 OTUs. Of these, 155 were identified as chimeras by UCHIME leaving 7,976 high quality OTUs. Representative sequences of these OTUs were then aligned using the QIIME implementation of PyNAST (Caporaso, Bittinger, et al., 2010) against the Greengenes core set (DeSantis et al., 2006). Of the remaining OTUs, 1,702 failed to align. These failed alignments were, invariably, co-amplified insect 16S and 18S, or chloroplast sequence. After filtering these OTUs, we were left with 7,355,166 sequences in 5,322 OTUs. The full OTU table is available as Table S4.

After all filtering and combining samples from individual colonies, we had at least 1,000 high-quality sequence reads from adults from 140 ant colonies and from larvae for 103 ant colonies. Rarefaction curves suggest that the diversity of these bacterial communities has been well-sampled by these data (Fig. S1). *Pseudomyrmex* bacterial communities are dominated by Proteobacteria, Firmictues, Bacteroidetes, and Actinobacteria (Fig. S2).

### Alpha diversity is higher in larvae

We measured alpha diversity as both the number of OTUs identified and using the Chao1 estimator in samples rarefied to 1,000 sequences. We tested for differences in alpha diversity between groups using Wilcoxon rank-sum tests and found that, for both metrics, alpha diversity was higher in larvae than adults in acacia-ants as a whole (*P. flavicornis, P. nigrocinctus*, and *P. spinicola* treated together), *P. gracilis*, and *P. spinicola* and for *Cephalotes* for the Chao1 estimator only (FDR-corrected P < 0.05, Fig. 2B). We did not find any significant difference between acacia-ants and *P. gracilis* using either metric (P > 0.10) or between countries for *P. gracilis* adults or larvae.

### Beta diversity in adults dominated by few genera

Principal coordinates analyses show clearly that *Cephalotes* and Dorylinae (army ants) dominate beta diversity in adult ants (Fig. 3A). *Camponotus* also groups quite tightly. All other taxa, including *Pseudomyrmex*, do not have apparent differences in beta diversity. We used machine learning to determine if automated classifiers could distinguish these genera. Indeed, the average error ratio of the 100 replicate classifiers for distinguishing *Cephalotes* from all other ants was 14.6 and for army ants was 2.2, though it failed for *Camponotus* (error ratio of 1.2). While clearly grouped in PCoA analyses, the *Camponotus* samples occupy the same coordinate space as many other genera, possibly explaining their lack of distinctiveness.

**Figure 3.**
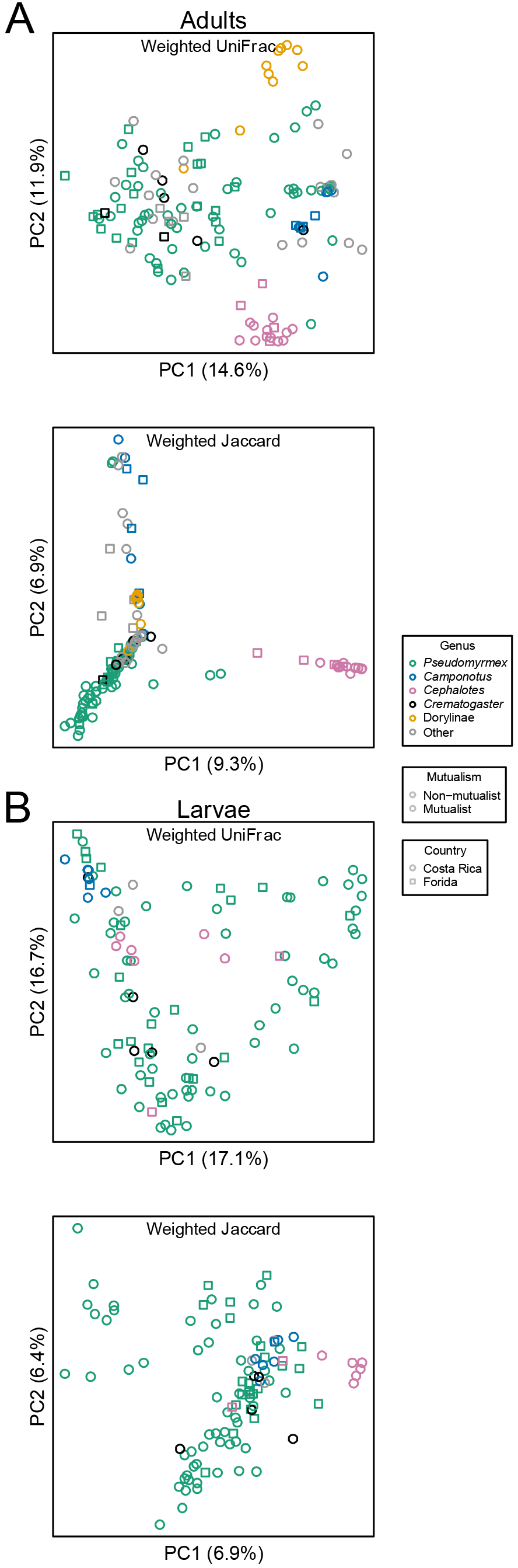
PCoA’s of weighted UniFrac and Jaccard beta diversities for adults (A) and larvae (B). Particularly for adults, *Cephalotes, Camponotus*, and Dorylinae (the army ants) clearly dominate differences in beta diversity.

We used Wilcoxon rank-sum tests to determine whether particular bacterial taxa differ in abundance between each of these three ant lineages and all other ants. We only tested the 255 OTUs that were present in at least 10 samples and used FDR-correction with the Benjamini-Hochberg procedure. There were four taxa that were significantly more abundant in army ants (FDR-corrected P < 1×10^−7^), all of which are classified as Tenericutes and three of which are classified as Entomoplasmataceae (OTUs 28, 2753, 2952). The fourth (OTU20) was not confidently classified below the level of phylum. All but OTU28 are ant-specific. The probability of drawing this many or more ant-specific taxa at random is 0.012.

There are 50 OTUs that are significantly more abundant in *Cephalotes* than other ants (FDR-corrected P < 0.05). Of these, 46 are classified as Proteobacteriea of which 12 are Alphaproteobacteria (10 are Rhizobiales and the rest are unclasssified), 18 are Betaproteobacteria (13 are Burkholderiales and the rest unclassified), and 12 are Gammaproteobacteria (two are Pseudomonadales and eight are Xanthomonadales with the last two unclassified). There are 35 ant-specific OTUs in this set. The probability of this many ant-specific OTUs is 2×10^−18^. There are also 10 OTUs that are significantly less abundant in *Cephalotes*, nine of which are classified as Proteobacteria. The last is Actinobacteria. Within the Proteobacteria, three are Rhizobiales, four are Burkholderiales, one is Pseudomonadales, and one is Xanthomonadales. None of these are ant-specific.

Although *Camponotus* was indistinguishable using supervised learning, five taxa were significantly more abundant in this genus than in all other ants (FDR-corrected P < 0.05). One of these is *Wolbachia*. The other four, as expected, are Enterobacteriaceae, one of which has no more detailed classification but the other three are all classified as *Blochmannia*. Four are ant-specific which has a probability of occurring at random of 0.0022.

### Larvae are less distinct

The same patterns hold generally true for between species comparisons of larvae, only to a much lesser extent. PCoA does show some grouping by genus (Fig. 3B) but supervised learning is unable to consistently distinguish between them. The error ratio for comparing *Cephalotes* larvae to other ant larvae is 1.9 and for *Camponotus*, it is essentially a random result with an error ratio of 1.05.

Again, the taxa that significantly differ in abundance between *Cephalotes* larvae and other ants are almost entirely restricted to the Proteobacteria. There are 25 OTUs that are significantly greater in abundance in *Cephalotes* larvae and 23 of them are Proteobacteria including six Rhizobiales (Alphaproteobacteria), four Burkholderiales (Betaproteobacteria), and eight Enterobacteriales (Gammaproteobacteria). The other two phyla represented are Bacteroidetes and Verrucomicrobia. Of these, 13 taxa distributed across the three classes are ant-specific. The probability of this many OTUs being ant-specific at random is 1×10^−5^. Lastly, only a single OTU classified as *Blochmannia* is significantly greater in abundance in *Camponotus* larvae.

### Geographic differences within species

We obtained samples for the species *P. gracilis* from both Florida and the ACG, making it ideal for assessing the impact of geographic location on bacterial communities without any external confounding influences. Supervised learning is unable to distinguish the two regions (error ratio of 1.19), none of the 166 OTUs present in at least four samples were significantly different in abundance after FDR correction (P > 0.5).

While the same patterns are apparent in the *P. gracilis* larvae from the two locations, distinguishing them is a bit more straightforward. The machine learning classifier had an error ratio of 1.43, still essentially a complete failure and no taxa are not significantly different in abundance after correcting for multiple testing.

However, differences are apparent from beta diversity analysis. We compared weighted Jaccard diversity within and between sites using nonparametric t-tests with Monte Carlo permutation. There is significantly lower beta diversity between all Florida samples than between all pairs of samples from Florida and the ACG (T = 2.7, P < 2×10^−16^). We find the same pattern when comparing within the ACG samples to all between country samples (T = 5.6, P < 2×10^−16^). The same pattern exists for larvae with significantly lower beta diversity in both those samples within Florida (T = 2.4, P = 0.002) and within the ACG (T = 3.9, P < 2×10^−16^) than those between countries. Although subtle, these differences precluded the direct comparison of bacterial communities from samples collected at the two distant sites.

### Small geographic distances cause little difference

Within Costa Rica we collected from four different sites, all within less than 10km of one another. We explored the potential for geographic influences on communities by individually examining those ant taxa for which we had collections from multiple sites including *Camponotus, Cephalotes, Crematogaster, P. gracilis, P. flavicornis, P. nigrocinctus*, and *P. spinicola*. We also combined all of the acacia ants (*P. flavicornis, P. nigrocinctus, P. spinicola*) for a single analysis. We found no signatures of consistent differences between samples collected from these different sites using supervised learning (Table S5).

For each of these ant taxa, we also tested for differences in beta diversity within sites and between sites using all four beta diversity metrics. Only a single test reaches significance for adults, that based on the unweighted Jaccard index for *P. gracilis* (T = 3.6, P = 0.010). However, in addition to *P. gracilis* larvae being significantly different using weighted Jaccard (P = 0.019), the acacia-ant larvae are significantly different using both of the unweighted metrics (T = 3.2, P = 0.02 for unweighted UniFrac and T = 3.5, P = 0.01 for unweighted Jaccard). It appears, therefore, that larvae may be more sensitive to small changes in environmental bacterial communities and, since only unweighted tests are significant, that only rare taxa change between sites.

### All species of acacia-ants are very similar

We see no indication that the three species of acacia-ants are distinct in any way. Tests of differences in beta diversity within and between species for all metrics were all non-significant (P > 0.2) for both gasters and larvae. Supervised learning has an error ratio of 0.95 for gasters and 1.07 for larvae.

### Larvae and adults are different

We examined differences between larvae and adults within species collected in the ACG. Machine learning classifiers perform quite well in these comparisons (Table S6). *Cephalotes, P. flavicornis, P gracilis*, and *P. spinicola* unequivocally have greater beta diversity between larvae and gasters than within those groups using any measure of beta diversity (FDR-corrected P < 0.01). The same is true when comparing all acacia-ants treated together. We don’t find any signature of this for *Camponotus, Crematogaster*, or *P. nigrocinctus*. However, these three taxa also have the lowest sample sizes so this may simply be a product of insufficient sampling.

For all within species tests of differences in abundance between larvae and adults, we only tested those OTUs present in at least four samples. Despite the lack of clear differences in beta diversity between *Crematogaster* life stages, there are three OTUs (none ant-specific) that are significantly more present in larvae than in adults as determined by a *χ*^2^ test (FDR-corrected P < 0.05) though not in abundance. All three of these are classified as Enterobacteriaceae. None of the 26 taxa tested were significantly different in *Camponotus*.

Within *Cephalotes*, we tested 130 taxa and 24 (16 are ant-specific: probability = 1×10^−8^) are more abundant in adults and 31 (3 ant-specific: probability = 0.86) are more abundant in larvae (FDR-corrected P < 0.05). Of those more abundant in gasters, 14 are Betaproteobacteria and 10 of these are Burkholderiales. Six are Gammaproteobacteria of which four are Xanthomonadales. In contrast, those more abundant in larvae are 11 Enterobacteriaceae, nine Rhizobiales, eight Lactobacillales, and two Actinomycetales.

Within *P. flavicornis*, we tested 302 taxa but none differ significantly in abundance. However, 19 (3 ant-specific: probability = 0.56) are significantly different in presence (χ^2^ test, FDR-corrected P < 0.05) and all of these are more often found in larvae. Of these, 10 are Actinomycetales and seven are Rhizobiales. We tested 198 taxa in *P. nigrocinctus* but none are significantly different in abundance or presence between larvae and adults.

*Pseudomyrmex spinicola* had 429 testable OTUs, four of which were significantly more abundant in adults (one Burkholderiales, one Rhizobiales, and two Acetobacteraceae (both of which are ant-specific: probability = 0.11)) and 102 of which were more abundant in larvae, only four of which are ant-specific (probability = 0.99). Of these, 57 are Alphaproteobacteria, 36 of which are Rhizobiales and 14 of which are Sphingomonadales. The majority of the rest (28) are Actinomycetales.

Of the 398 taxa tested for *P. gracilis*, none are significantly more abundant in gasters and seven are more abundant in larvae, including four Actinomycetales and three Rhizobiales. Only one is ant-specific (probability = 0.68).

Lastly, we combined the three acacia-ants to test 829 OTUs for differences between gasters and larvae. While 14 are more abundant in adults (only four are ant-specific: probability = 0.15), 131 are more abundant in larvae (FDR-corrected P < 0.05), of which only 12 are ant-specific (probability = 0.98). All of these significantly different taxa are similarly distributed taxonomically as for the other comparisons. For those that are more abundant in adults, three are Actinomycetales, three are Rhizobiales, and six are Acetobacteraceae. For those more abundant in larvae, 49 are Rhizobiales, 39 are Actinomycetales, 11 are Sphingomonadales, five are Rickettsiales, five are Acetobacteraceae, seven are Solirubrobacterales, and five are Lactobacillales.

### Mutualists versus generalists

We compared acacia-ants to the non-mutualistic but sympatric species *P. gracilis* using only samples from Costa Rica to determine if there is anything unique about the acacia-ant microbiome. We also tested acacia-ant communities against the *Crematogaster* samples that were found parasitizing acacia trees.

Supervised learning was unable to distinguish *P. gracilis* from acacia-ants with an error ratio of 1.46 for gasters and 1.19 for larvae. Similarly, the comparison with *Crematogaster* achieved an error ratio of 1.19 for larvae and 1.14 for gasters. However, comparisons of all beta diversity metrics within acacia-ants and all beta diversities between acacia-ants and either *P. gracilis* or *Crematogaster* are all significant (FDR-corrected P < 0.001) for both adults and larvae.

We tested those OTUs present in at least 10 samples for differences in abundance between *Pseudomyrmex* acacia-ants and *P. gracilis*. Of the 95 bacterial taxa tested, five are significantly more abundant in acacia-ants than in *P. gracilis* adults, four of which are classified as Acetobacteraceae (OTU5124, OTU3420, OTU4488 and OTU84; Wilcoxon rank-sum FDR-corrected P < 0.05) and the last of which is Cytophagaceae (OTU44). This last taxon has a 16S rRNA sequence that is 100% similar to sequence obtained from the human skin microbiome (KF063112). OTU44 is, therefore, likely to be a widespread commensal or present as the result of latent contamination from the environment or laboratory. The four Acetobacteraceae taxa are ant-specific (probability = 0.0022), but are present in both acacia-ants and *P. gracilis* so they are not unique to mutualists. However, both OTU84 and OTU44 make up substantial quantities in mutualists with an average of 9.8% and 7.6% of sequences, respectively and less than 1% in *P. gracilis* (Fig. 4). All of the other significant taxa are present at less than 1% frequency. For larvae, 353 OTUs were tested and one was significantly different in abundance between acacia-ants and *P. gracilis* (OTU19: Actinomycetales). This OTU is also ant-specific and makes up an average of 6.3% of sequences in mutualists and less than 0.01% of sequences in *P. gracilis*. We also combined all OTUs from individual bacterial families and tested for differences between mutualists and *P. gracilis* but only Cytophagaceae was significantly different. Lastly, we tested for differences in family abundance in larvae but none were significantly different (FDR-corrected P > 0.05).

**Figure 4.**
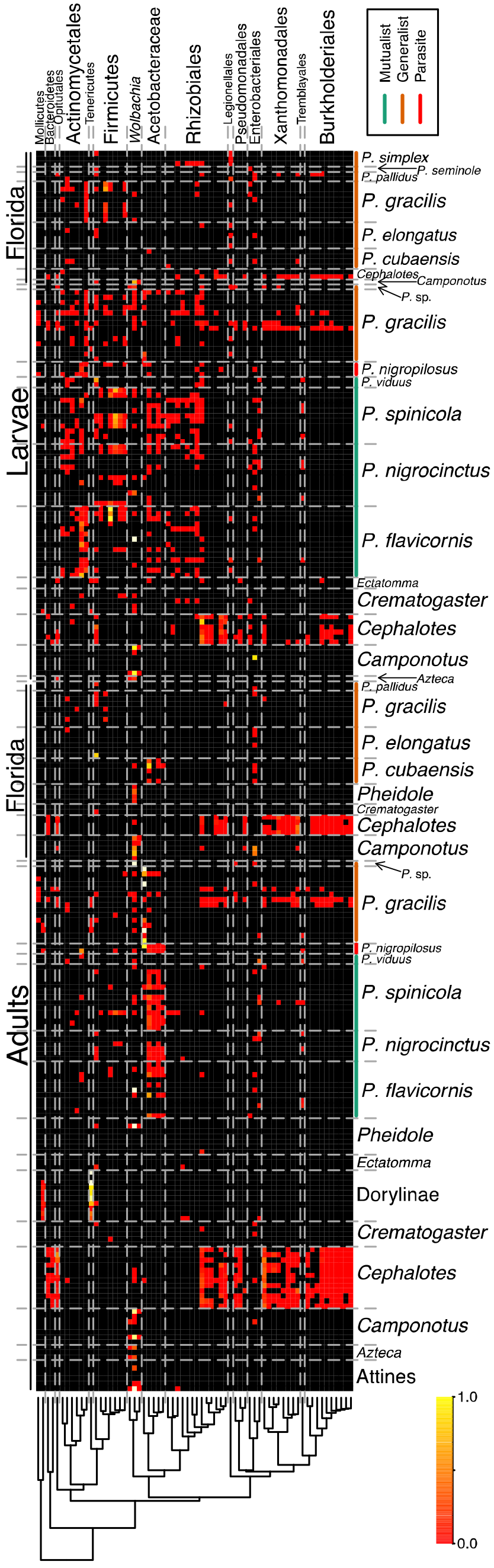
Heatmap of ant-specific OTUs that occur in at least 10 samples. Any occurrence within a sample of fewer than five reads was discarded as likely cross-sample contamination. Colors represent the proportion of all reads from each individual sample. Acacia-ants are indicated by the green lines and the generalist that they were compared to (*P. gracilis*) is indicated by the orange line. *P. viduus* is a mutualist with another type of plant, *Triplaris* and *P. nigropilosus* is an obligate parasite of acacia-plants.

For *Crematogaster* we tested 54 OTUs in adults and 246 in larvae but none were significantly different in abundance after multiple-test correction. We, therefore, tested for differences in presence using *χ*^2^ tests and found that OTU44, the Cytophagaceae cited above, is significantly more frequently encountered in acacia-ants (FDR-corrected P = 0.043). Before correction, 10 taxa are significantly more abundant in mutualists than in *Crematogaster* including all of the taxa identified in the comparison with *P. gracilis* (P < 0.05) as well as a Rhizobiales (OTU11) and another Acetobacteraceae (OTU9).

Most of the ant-specific taxa found in higher levels in gasters or larvae of acacia-ants are also found in other ants to varying degrees. For OTU19, there are a total of 10 reads found in non-*Pseudomyrmex* species, for OTU84 there are 67, OTU5124 there are three and for OTU4488 there is one. For OTU36, a relative of OTU19, there are 221. There are none for OTU3420. Given these small numbers, it is possible that these taxa are actually *Pseudomyrmex* specific and that between sample contamination or barcode errors are causing the appearance of these taxa in other species.

### Similarity to previously identified taxa

The clade of ant-specific OTUs in the Acetobacteraceae that differ in abundance in mutualists are of particular interest (Fig. 5). OTU84 is 97% similar to Acetobacteraceae from bee guts (KU572006) and OTU5124, OTU4488 and OTU3420 are all 98% similar to a sequence obtained from the Argentine ant, *Linepithema humile* (KX984918). OTU9 is, in contrast, 100% similar to previously sequenced *Asaia* isolated from a variety of insects.

**Figure 5.**
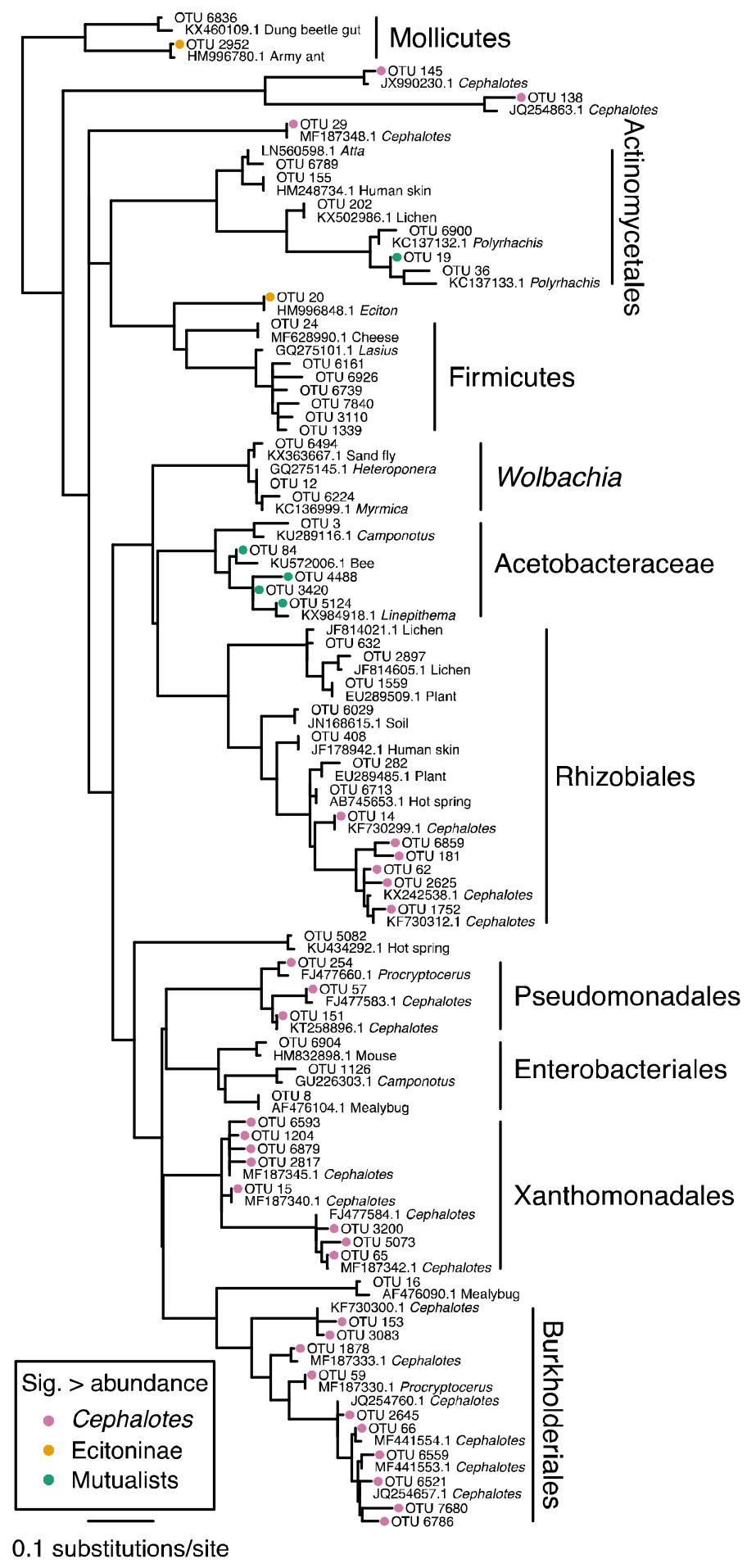
Phylogeny of ant-specific OTUs that occur in at least 10 samples and their closest hits in NCBI’s nucleotide database. Colored circles indicate those taxa that are significantly more abundant in *Cephalotes* compared with all other ants, Dorylinae compared with all other ants, and mutualistic *Pseudomyrmex* compared with non-mutualistic *Pseudomyrmex*.

The Actinomycetales identified as different in abundance between mutualists and non-mutualists are also potentially influencing the ecology of these species (Fig. 5). OTU19 is 96% similar to a sequence from *Polyrhachis* (KC137132) and OTU36 is 93% similar to another sequence from *Polyrhachis* (KC137133). The two OTU’s are 95% divergent to each other.

### Ant-specific analyses increase accuracy

Limiting the OTUs input into supervised learning analyses to just those found to be ant-specific increases their accuracy. Note that the decrease in read numbers meant that we had to rarefy the OTU table to 500 rather than 1000. Between *P. gracilis* and acacia-ants, the error ratio was 2.35 for adults, though it was still only 1.03 for larvae. For the comparison between *Crematogaster* and acacia-ants, the error ratio remained low at 1.39. For the comparison of *P. gracilis* between countries, the gasters have an improved error ratio of 1.61, though it is still not significantly distinguishable. For comparisons between ant genera, the error ratio for *Camponotus* is 1.5, 7.53 for *Cephalotes*, and 8.5 for *Eciton*.

### Antibiotic genes present

There were 8,926,317 reads with discernible taxonomy in the *P. nigropilosus* larval metagenome. Of these, the vast majority were classified as Proteobacteria (8,200,012). The second most abundant phylum was Firmicutes with 588,226 reads and the third was Actinobacteria with 33,578. Of the Actinobacteria, the vast majority were Actinomycetales (29,920). The Proteobacteria are likely the result of an infection in the individuals sequenced as 7,395,038 reads were classified as Enterobacteriaceae yet only 573,405 of all reads in the 16S rDNA amplicon dataset and only 57 of the amplicon reads from the same colony (BER0554) were classified to this family.

In the *P. flavicornis* adult metagenome there were 2,517,613 taxonomic hits and 2,292,522 of these were to Proteobacteria, 35,103 were to Firmictues and 27,047 were to Actinobacteria. However, in this case, 1,662,817 of the Proteobacteria reads were to Acetobacteraceae.

Of the hits to Actinomycetales, 8,777 were assigned SEED subsystem function in *P. flavicornis* adults and 6,389 in *P. nigropilosus* larvae. In larvae, 533 are assigned to secondary metabolism and 503 of these are classified as non-ribosomal peptide synthetase genes which are likely to be involved in the production of defensive chemicals. In contrast, only four reads are assigned to secondary metabolism among the Actinobacteria reads in *P. flavicornis* adults, none of which are related to antibiotic production. Although it was not possible to associate these genes with particular OTUs, the *P. nigropilosus* larvae from the same colony as the individuals for which we sequenced metagenomes yielded a total of 6,307 Actinomycetales 16S reads and 1,336 (21%) of these were derived from OTU19 and 1,498 (24%) were from OTU36. The third most abundant OTU was OTU7467 which was represented by 1,066 (17%) reads. All other Actinomycetales OTUs were represented by fewer than 400 reads. Therefore, the Actinomycetales genes detected in the *P. nigropilosus* metagenome are likely from one of these three OTUs, two of which (OTU19 and OTU36) are ant-specific and distantly related to other taxa (Fig. 5).

Within *P. flavicornis* adults, there are 760,202 functional hits for Acetobacteraceae, 4,017 of which are to nitrogen metabolism. Of these, 4,014 are involved in ammonia assimilation and the rest are classified as nitrate and nitrite ammonification. More than half (2,049) are classified as type 1 glutamine synthetase (EC 6.3.1.2) and most of the rest (1,717) are glutamate synthase (EC 1.4.1.13). The presence of these functional groups is not particularly noteworthy as they are present throughout the bacterial phylogeny. No genes involved in nitrogen fixation were detected from the Acetobacteraceae. Only 3,844 reads are Acetobacteraceae in *P. nigropilosus* larvae which yielded 1,124 functional hits. Of these, 64 are to nitrogen metabolism. Three of these are to ammonia assimilation and 61 are to nitrate and nitrite ammonification of which 56 are either nitrate or nitrate reductases. Overall, no evidence for protein enrichment is apparent from our metagenomic data.

## Discussion

### Bacteria are not essential to ant herbivory

Several clades of ants have clearly coevolved with specific clades of bacteria and, at least for the partnership between *Camponotus* carpenter ants and their *Blochmannia* symbionts, these relationships do yield nutritional benefits (de Souza, Bézier, Depoix, Drezen, & Lenoir, 2009; Feldhaar et al., 2007). Recent work in the ant genus *Cephalotes*, a strict herbivore, has shown the presence of coevolved Rhizobiales bacteria potentially enriching the protein-deficient diets of their hosts (Hu et al., 2014; Russell et al., 2009; Sanders et al., 2014). Tempting though it may be to use these taxa to draw broader conclusions about the necessity of gut symbionts for the evolution of herbivory, it now appears that these relationships are exceptional among ants (Russell, Sanders, & Moreau, 2017). Notably, while large numbers of bacteria are not necessarily required for symbiotic relationships with their hosts, those ants known to engage in close relationships with bacteria, *Cephalotes*, *Camponotus*, and army ants (Funaro et al., 2011; Łukasik et al., 2017), host the highest numbers of bacteria, well above the numbers present in acacia-ants and ants generally (Fig. 2; (Sanders et al., 2017)). These taxa also dominate beta diversity, further supporting their exceptional nature (Fig. 3).

Acacia-ants in the genus *Pseudomyrmex* are among the strictest of insect herbivores (Fig. 1), yet they do not host specific coevolved bacteria, and are likely to obtain complete nutrition from their host plants, as has been previously suggested (Heil, Kruger, Baumann, & Linsenmair, 2004). Instead, the taxa in their guts are spread across *Pseudomyrmex* species with varied diets and similar taxa are present in distantly related ants and insects more generally. However, the relative abundance of these taxa differs between *Pseudomyrmex* mutualists with limited diets and generalists with more diverse and protein-rich diets. Specifically, there are two clades of bacteria which differ in quantity between mutualists ant non-mutualists, Acetobacteraceae and Actinomycetales.

### Acetobacteraceae

Acetic acid bacteria are common inhabitants of insect guts, particularly among those that feed largely on nectar (Crotti et al., 2010, 2016) and including two widespread groups of ants, *Camponotus* (Brown & Wernegreen, 2016) and *Linepithema* (Hu et al., 2017). These taxa play a role in larval survival in honey bees (Corby-Harris et al., 2014), larval development time in *Anopheles* (Chouaia et al., 2012), and development and metabolism in *Drosophila* (Shin et al., 2011). While, to our knowledge, there are no reports of species-specific relationships between hosts and acetic acid bacteria, the genomes of several lineages inhabiting honey bees and mosquitoes indicate that adaptations for insect association have occurred (Chouaia et al., 2014).

It is difficult to speculate on the cause or consequence of the differences in Acetobacteraceae abundance between the acacia mutualists and non-mutualists. Though closely related and likely performing largely similar roles, diverse strains of bumble bee associated Acetobacteraceae have been found to have diverse functions (Engel, Martinson, & Moran, 2012; Powell, Leonard, Kwong, Engel, & Moran, 2016). It is, therefore, quite possible that the different strains of acetic acid bacteria in *Pseudomyrmex* are influencing their hosts in different ways, though it is also possible that these bacteria are simply colonizing the environments to which they are most suited; environments that likely differ predominantly in the relative amount of sugar. Regardless, it is obvious that Acetobacteraceae are a core member of the gut bacteria present in a wide diversity of ants and insects (Crotti et al., 2010, 2016; Kautz, Rubin, & Moreau, 2013).

### Actinomycetales

Though there are a few examples of Actinomycetales likely providing nutritional assistance to their insect hosts (Ben-Yakir, 1987; Durvasula et al., 2008; Haas & König, 1988), defensive mutualisms are much more common (Kaltenpoth, 2009). The leaf-cutter ants engage in a well-studied defensive mutualism with bacteria in the genus *Pseudonocardia*, growing these bacteria on specific locations on their cuticles and using the compounds produced to eliminate a parasitic fungus from their gardens (Currie, Poulsen, Mendenhall, Boomsma, & Billen, 2006; Currie, Scott, Summerbell, & Malloch, 1999; Haeder, Wirth, Herz, & Spiteller, 2009). Another group of obligate plant-ants, *Allomerus*, also host Actinobacteria that they seem to use in a similar way, selecting for just one fungal lineage within their nests (Ruiz-Gonzalez et al., 2011; Seipke et al., 2012). And several other obligate plant-nesting ant species are known to host related lineages, including another acacia-ant (Hanshew et al., 2015). Although associations with fungus have not been reported in *Pseudomyrmex*, Actinomycetales clearly makes up a core component of the *Pseudomyrmex* microbiome and includes antibiotic biosynthesis genes. Given the much greater abundance of Actinomycetales taxa in larvae than adults, these bacteria are likely helping to defend juveniles from pathogens as occurs in beewolf wasps (Kaltenpoth, Göttler, Herzner, & Strohm, 2005). As yet, it is not clear why *Pseudomyrmex* in particular would require such protections or how this relationship may differ in mutualists and non-mutualists.

### Adults and larvae host very different bacteria

Outside of the specific bacterial taxa associated with *Pseudomyrmex*, we also make several findings regarding ant-microbe interactions more generally. First, bacterial communities are drastically different in adults and larvae, a pattern especially apparent in our *Pseudomyrmex* samples (Fig. 4). The Acetobacteraceae OTUs are only significantly more abundant in mutualists in adults and Actinomycetales are, in contrast, only present in abundance and significantly different in larvae. Focusing on just one of either of these life stages would have led to the failure to identify these taxa as important community members.

The higher alpha diversities within larvae were quite unexpected. However, these may be due simply to the limitations of larval physiology. Ant larvae have closed digestive systems, thus, any bacteria present in their food may remain in their guts until pupation. The difference with adults suggests that environmental contaminants in adult food may be transient enough as to be uninfluential. Many of those taxa that differed in abundance between larvae and adults are not ant-specific but instead appeared to be widespread environmental bacteria.

### Ant-specific bacteria differ across species

Finally, the bacterial taxa that differ in abundance between ant genera and behaviors tend to be those that associate specifically with ants rather than those that are present in any quantity large enough to be detected from the environment more generally (Fig. 5). These bacterial lineages may have adaptations for occupying ants or insects more generally, and may, in some cases, coevolve to take advantage of the insect gut environment, despite not being tied to a specific host. The relatively low quantities of bacteria found in many distantly related types of ants and the frequent dominance of those communities by acetic acid bacteria, as we find for *Pseudomyrmex*, supports this hypothesis (Hu et al., 2017; Russell et al., 2017; Sanders et al., 2017).

## Conclusions

Although the acacia-ant diet may be richer than other ant herbivores thanks to the mutualistic relationship with their host plants, we find no support for the hypothesis that bacterial partners are required for the evolution of herbivory. Several other lines of evidence pointing in this direction have also accumulated recently, including in other groups of ants with herbivorous diets (Hu et al., 2017) and even in caterpillars which appear to host no bacteria at all (Hammer, Janzen, Hallwachs, Jaffe, & Fierer, 2017). Instead, defensive symbiosis with Actinobacteria, an as yet unexplored aspect of the acacia-ant mutualism, appears quite likely.

## Acknowledgements

For assistance in the field and lab we thank Elizabeth Pringle, Arista Tischner, Alexandra Westrich, and Max Winston. For helpful conversations that improved this study we thank Jack Gilbert, Winnie Hallwachs, Daniel Janzen, Jacob Russell, Jon Sanders, and members of the Moreau lab. For permits to collect specimens in Costa Rica we thank Daniel Janzen and the Area de Conservación Guanacaste. In the Florida Keys we thank the United States Fish and Wildlife Service, Florida Department of Environmental Protection, and the Nature Conservancy for permission to collect specimens. This project was made possible by financial support from the National Science Foundation (NSF DDIG DEB-1311417 to B.E.R.R and C.S.M. and NSF DEB-1050243 to C.S.M.).

---

**Figure S1.**
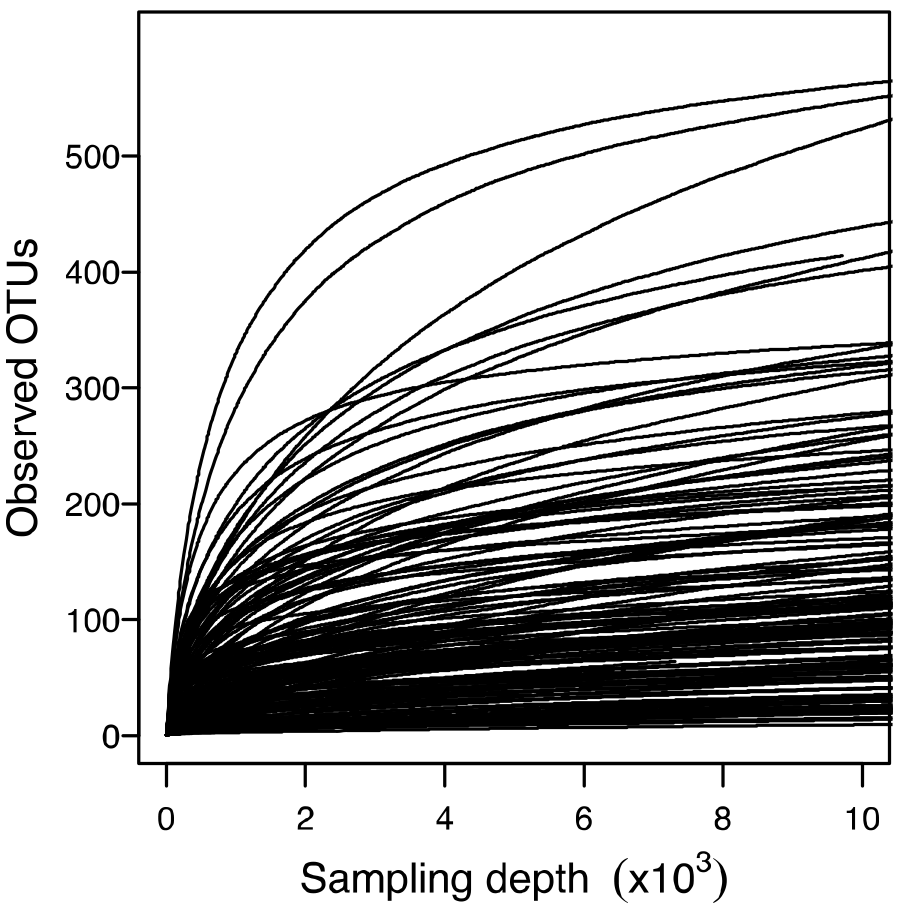
Rarefaction plots for all samples.

**Figure S2.**
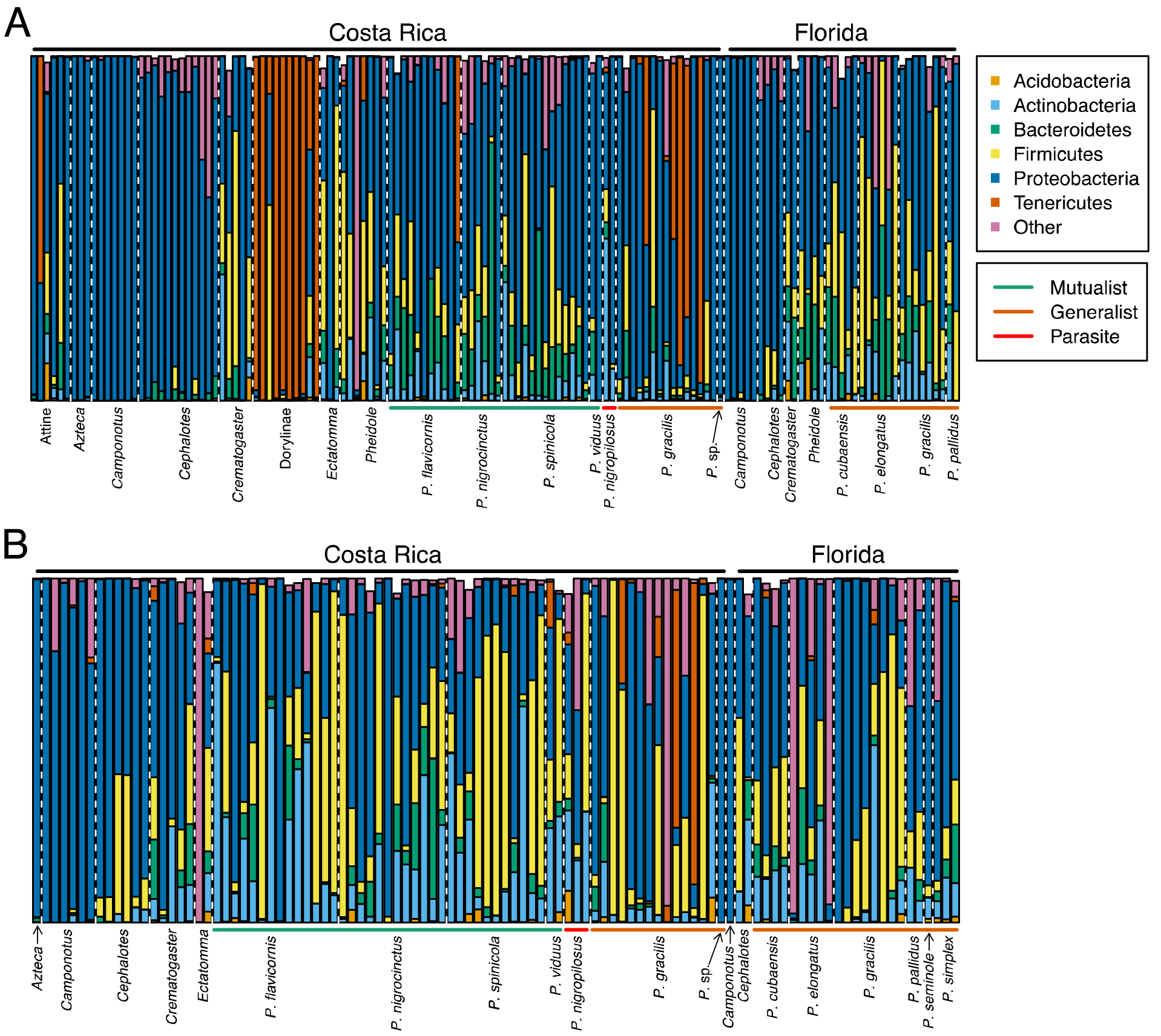
Barplots showing relative abundances of bacterial phyla.

## Supplementary Tables

Table S1. Details for all samples used in this study.

Table S2. Wilcoxon tests for differences in nitrogen isotope ratios.

Table S3. Tests for differences in bacterial quantities.

Table S4. OTU table.

Table S5. Supervised learning analyses between sites in ACG.

Table S6. Supervised learning analyses between adults and larvae.

## References

Bandi, C., Sironi, M., Damiani, G., Magrassi, L., Nalepa, C. A., Laudani, U., & Sacchi, L. (1995). The establishment of intracellular symbiosis in an ancestor of cockroaches and termites. Proceedings. Biological Sciences, 259(1356), 293–299. doi:10.1098/rspb.1995.0043

Benemann, J. R. (1973). Nitrogen fixation in termites. Science, 181(4095), 164–165.

Ben-Yakir, D. (1987). Growth retardation of Rhodnius prolixus symbionts by immunizing host against Nocardia (Rhodococcus) rhodnii. Journal of Insect Physiology, 33(6), 379–383.

Bolger, A. M., Lohse, M., & Usadel, B. (2014). Trimmomatic: a flexible trimmer for Illumina sequence data. Bioinformatics, 30(15), 2114–2120. doi:10.1093/bioinformatics/btu170

Borm, S. v., Buschinger, A., Boomsma, J. J., & Billen, J. (2002). Tetraponera ants have gut symbionts related to nitrogen-fixing root-nodule bacteria. Proceedings of the Royal Society B: Biological Sciences, 269(1504), 2023–2027. doi:10.1098/rspb.2002.2101

Breznak, J. A., Winston, J. B., Mertins, J. W., & Coppel, H. C. (1973). Nitrogen fixation in termites. Nature, 244, 577–580.

Brown, B. P., & Wernegreen, J. J. (2016). Deep divergence and rapid evolutionary rates in gut-associated Acetobacteraceae of ants. BMC Microbiology, 16(140), 140. doi:10.1186/s12866-016-0721-8

Caporaso, J. G., Bittinger, K., Bushman, F. D., DeSantis, T. Z., Andersen, G. L., & Knight, R. (2010). PyNAST: a flexible tool for aligning sequences to a template alignment. Bioinformatics, 26(2), 266–267. doi:10.1093/bioinformatics/btp636

Caporaso, J. G., Kuczynski, J., Stombaugh, J., Bittinger, K., Bushman, F. D., Costello, E. K., … Knight, R. (2010). QIIME allows analysis of high-throughput community sequencing data. Nature Methods, 7(5), 335–336.

Caporaso, J. G., Lauber, C. L., Walters, W. A., Berg-Lyons, D., Huntley, J., Fierer, N., … Bauer, M. (2012). Ultra-high-throughput microbial community analysis on the Illumina HiSeq and MiSeq platforms. The ISME Journal, 6(8), 1621–1624.

Cheng, Q., & Aksoy, S. (1999). Tissue tropism, transmission and expression of foreign genes in vivo> in midgut symbionts of tsetse flies. Insect Molecular Biology, 8(1), 125–132.

Chomicki, G., Ward, P. S., & Renner, S. S. (2015). Macroevolutionary assembly of ant/plant symbioses: Pseudomyrmex ants and their ant-housing plants in the Neotropics. Proceedings of the Royal Society B: Biological Sciences, 282(1819), 20152200. doi:10.1098/rspb.2015.2200

Chouaia, B., Gaiarsa, S., Crotti, E., Comandatore, F., Degli Esposti, M., Ricci, I., … Daffonchio, D. (2014). Acetic acid bacteria genomes reveal functional traits for adaptation to life in insect guts. Genome Biology and Evolution, 6(4), 912–920. doi:10.1093/gbe/evu062

Chouaia, B., Rossi, P., Epis, S., Mosca, M., Ricci, I., Damiani, C., … others. (2012). Delayed larval development in Anopheles mosquitoes deprived of Asaia bacterial symbionts. BMC Microbiology, 12(1), S2.

Clement, L. W., Köppen, S. C. W., Brand, W. A., & Heil, M. (2008). Strategies of a parasite of the ant-acacia mutualism. Behavioral Ecology and Sociobiology, 62(6), 953–962. doi:10.1007/s00265-007-0520-1

Corby-Harris, V., Snyder, L. A., Schwan, M. R., Maes, P., McFrederick, Q. S., & Anderson, K. E. (2014). Origin and effect of Alpha 2.2 Acetobacteraceae in honey bee larvae and description of Parasaccharibacter apium gen. nov., sp. nov. Applied and Environmental Microbiology 80(24), 7460–7472. doi:10.1128/AEM.02043-14

Crotti, E., Chouaia, B., Alma, A., Favia, G., Bandi, C., Bourtzis, K., & Daffonchio, D. (2016). Acetic acid bacteria as symbionts of insects. In K. Matsushita, H. Toyama, N. Tonouchi, & A. Okamoto-Kainuma (Eds.), Acetic Acid Bacteria: Ecology and Physiology (pp. 121–142). Tokyo: Springer Japan. doi:10.1007/978-4-431-55933-7_5

Crotti, E., Rizzi, A., Chouaia, B., Ricci, I., Favia, G., Alma, A., … Daffonchio, D. (2010). Acetic acid bacteria, newly emerging symbionts of insects. Applied and Environmental Microbiology 76(21), 6963–6970. doi:10.1128/AEM.01336-10

Currie, C. R., Poulsen, M., Mendenhall, J., Boomsma, J. J., & Billen, J. (2006). Coevolved crypts and exocrine glands support mutualistic bacteria in fungus-growing ants. Science, 311(5757), 81–83.

Currie, C. R., Scott, J. A., Summerbell, R. C., & Malloch, D. (1999). Fungus-growing ants use antibiotic-producing bacteria to control garden parasites. Nature, 398(6729), 701.

Davidson, D. W. (1997). The role of resource imbalances in the evolutionary ecology of tropical arboreal ants. Biological Journal of the Linnean Society, 61(2), 153–181. doi:10.1111/j.1095-8312.1997.tb01785.x

Davidson, D. W., Cook, S. C., Snelling, R. R., & Chua, T. H. (2003). Explaining the abundance of ants in lowland tropical rainforest canopies. Science, 300, 969–972.

Davidson, D. W., & Patrell-Kim, L. (1996). Tropical arboreal ants: Why so abundant? In A. C. Gibson (Ed.), Neotropical Biodiversity and Conservation (pp. 127–140). Los Angeles, CA: UCLA Botanical Garden.

de Souza, D. J., Bézier, A., Depoix, D., Drezen, J.-M., & Lenoir, A. (2009). Blochmannia endosymbionts improve colony growth and immune defence in the ant Camponotus fellah. BMC Microbiology, 9(1), 29. doi:10.1186/1471-2180-9-29

DeSantis, T. Z., Hugenholtz, P., Larsen, N., Rojas, M., Brodie, E. L., Keller, K., … Andersen, G. L. (2006). Greengenes, a chimera-checked 16S rRNA gene database and workbench compatible with ARB. Applied and Environmental Microbiology, 72(7), 5069–5072. doi:10.1128/AEM.03006-05

Durvasula, R. V., Sundaram, R. K., Kirsch, P., Hurwitz, I., Crawford, C. V., Dotson, E., & Beard, C. B. (2008). Genetic transformation of a Corynebacterial symbiont from the Chagas disease vector Triatoma infestans. Experimental Parasitology, 119(1), 94–98. doi:10.1016/j.exppara.2007.12.020

Edgar, R. C. (2013). UPARSE: highly accurate OTU sequences from microbial amplicon reads. Nature Methods, 10, 996–998. doi:10.1038/nmeth.2604

Eilmus, S., & Heil, M. (2009). Bacterial associates of arboreal ants and their putative functions in an obligate ant-plant mutualism. Applied and Environmental Microbiology, 75(13), 4324–4332. doi:10.1128/AEM.00455-09

Engel, P., Martinson, V. G., & Moran, N. A. (2012). Functional diversity within the simple gut microbiota of the honey bee. Proceedings of the National Academy of Sciences, 109(27), 11002–11007. doi:10.1073/pnas.1202970109

Feldhaar, H., Straka, J., Krischke, M., Berthold, K., Stoll, S., Mueller, M. J., & Gross, R. (2007). Nutritional upgrading for omnivorous carpenter ants by the endosymbiont Blochmannia. BMC Biology, 5(1), 48. doi:10.1186/1741-7007-5-48

Flint, H. J., Bayer, E. A., Rincon, M. T., Lamed, R., & White, B. A. (2008). Polysaccharide utilization by gut bacteria: potential for new insights from genomic analysis. Nature Reviews Microbiology, 6(2), 121–131. doi:10.1038/nrmicro1817

Floren, A., Biun, A., & Linsenmair, E. K. (2002). Arboreal ants as key predators in tropical lowland rainforest trees. Oecologia, 131(1), 137–144. doi:10.1007/s00442-002-0874-z

Funaro, C. F., Kronauer, D. J. C., Moreau, C. S., Goldman-Huertas, B., Pierce, N. E., & Russell, J. A. (2011). Army ants harbor a host-specific clade of Entomoplasmatales bacteria. Applied and Environmental Microbiology, 77(1), 346–350. doi:10.1128/AEM.01896-10

Haas, F., & König, H. (1988). Coriobacterium glomerans gen. nov., sp. nov. from the intestinal tract of the red soldier bug. International Journal of Systematic and Evolutionary Microbiology, 38(4), 382–384.

Haeder, S., Wirth, R., Herz, H., & Spiteller, D. (2009). Candicidin-producing Streptomyces support leaf-cutting ants to protect their fungus garden against the pathogenic fungus Escovopsis. Proceedings of the National Academy of Sciences, 106(12), 4742–4746.

Hammer, T. J., Janzen, D. H., Hallwachs, W., Jaffe, S. P., & Fierer, N. (2017). Caterpillars lack a resident gut microbiome. Proceedings of the National Academy of Sciences, 114(36), 9641–9646. doi:10.1073/pnas.1707186114

Hansen, A. K., & Moran, N. A. (2011). Aphid genome expression reveals host-symbiont cooperation in the production of amino acids. Proceedings of the National Academy of Sciences, 108(7), 2849–2854. doi:10.1073/pnas.1013465108

Hanshew, A. S., McDonald, B. R., Díaz Díaz, C., Djiéto-Lordon, C., Blatrix, R., & Currie, C. R. (2015). Characterization of Actinobacteria associated with three ant-plant mutualisms. Microbial Ecology 69(1), 192–203. doi:10.1007/s00248-014-0469-3

Heil, M., Kruger, R., Baumann, B., & Linsenmair, E. K. (2004). Main nutrient compounds in food bodies of Mexican acacia ant-plants. Chemoecology, 14(1), 45–52. doi:10.1007/s00049-003-0257-x

Hu, Y., Holway, D. A., Łukasik, P., Chau, L., Kay, A. D., LeBrun, E. G., … Russell, J. A. (2017). By their own devices: invasive Argentine ants have shifted diet without clear aid from symbiotic microbes. Molecular Ecology, 26(6), 1608–1630. doi:10.1111/mec.13991

Hu, Y., Łukasik, P., Moreau, C. S., & Russell, J. A. (2014). Correlates of gut community composition across an ant species (Cephalotes varians) elucidate causes and consequences of symbiotic variability. Molecular Ecology, 23(6), 1284–1300. doi:10.1111/mec.12607

International Aphid Genomics Consortium. (2010). Genome sequence of the pea aphid Acyrthosiphon pisum. PLoS Biology, 8(2), e1000313.

Janzen, D. H. (1966). Coevolution of mutualism between ants and acacias in Central America. Evolution, 20(3), 249–279. doi:10.2307/2406628

Janzen, D. H. (1967). Interaction of the bull's-horn acacia (Acacia cornigera L.) with an ant inhabitant (Pseudomyrmex ferruginea F. Smith) in eastern Mexico. Kansas University Scientific Bulletin, 47, 315–558.

Janzen, D. H. (1975). Pseudomyrmex nigropilosa: A parasite of a mutualism. Science, 188(4191), 936–937.

Kaltenpoth, M. (2009). Actinobacteria as mutualists: general healthcare for insects? Trends in Microbiology, 17(12), 529–535. doi:10.1016/j.tim.2009.09.006

Kaltenpoth, M., Göttler, W., Herzner, G., & Strohm, E. (2005). Symbiotic bacteria protect wasp larvae from fungal infestation. Current Biology, 15(5), 475–479.

Kautz, S., Rubin, B. E. R., & Moreau, C. S. (2013). Bacterial infections across the ants: Frequency and prevalence of Wolbachia, Spiroplasma, and Asaia. Psyche: A Journal of Entomology, 2013, 1–11. doi:10.1155/2013/936341

Łukasik, P., Newton, J. A., Sanders, J. G., Hu, Y., Moreau, C. S., Kronauer, D. J. C., … Russell, J. A. (2017). The structured diversity of specialized gut symbionts of the New World army ants. Molecular Ecology, 26(14), 3808–3825. doi:10.1111/mec.14140

Lunter, G., & Goodson, M. (2011). Stampy: A statistical algorithm for sensitive and fast mapping of Illumina sequence reads. Genome Research, 21(6), 936–939. doi:10.1101/gr.111120.110

Majer, J. D. (1990). The abundance and diversity of arboreal ants in northern Australia. Biotropica, 22(2), 191. doi:10.2307/2388412

Miura, T., Braendle, C., Shingleton, A., Sisk, G., Kambhampati, S., & Stern, D. L. (2003). A comparison of parthenogenetic and sexual embryogenesis of the pea aphid Acyrthosiphon pisum (Hemiptera: Aphidoidea). Journal of Experimental Zoology Part B: Molecular and Developmental Evolution, 295(1), 59–81.

Moran, N. A., & Baumann, P. (2000). Bacterial endosymbionts in animals. Current Opinion in Microbiology, 3(3), 270–275.

Moreau, C. S., & Rubin, B. E. R. (2017). Diversity and persistence of the gut microbiome of the giant neotropical bullet ant. Integrative and Comparative Biology, 57(4), 682–689. doi:10.1093/icb/icx037

Navas-Molina, J. A., Peralta-Sánchez, J. M., González, A., McMurdie, P. J., Vázquez-Baeza, Y., Xu, Z., … Knight, R. (2013). Advancing our understanding of the human microbiome using QIIME. In Methods in Enzymology (Vol. 531, pp. 371–444). Elsevier. doi:10.1016/B978-0-12-407863-5.00019-8

Powell, J. E., Leonard, S. P., Kwong, W. K., Engel, P., & Moran, N. A. (2016). Genome-wide screen identifies host colonization determinants in a bacterial gut symbiont. Proceedings of the National Academy of Sciences, 113(48), 13887–13892. doi:10.1073/pnas.1610856113

Ramalho, M. O., Bueno, O. C., & Moreau, C. S. (2017). Microbial composition of spiny ants (Hymenoptera: Formicidae: Polyrhachis) across their geographic range. BMC Evolutionary Biology, 17(1). doi:10.1186/s12862-017-0945-8

Rubin, B. E. R., & Moreau, C. S. (2016). Comparative genomics reveals convergent rates of evolution in ant-plant mutualisms. Nature Communications, 7, 12679. doi:10.1038/ncomms12679

Rubin, B. E. R., Sanders, J. G., Hampton-Marcell, J., Owens, S. M., Gilbert, J. A., & Moreau, C. S. (2014). DNA extraction protocols cause differences in 16S rRNA amplicon sequencing efficiency but not in community profile composition or structure. MicrobiologyOpen, 3(6), 910–921. doi:10.1002/mbo3.216

Ruiz-Gonzalez, M. X., Male, P.-J. G., Leroy, C., Dejean, A., Gryta, H., Jargeat, P., … Orivel, J. (2011). Specific, non-nutritional association between an ascomycete fungus and Allomerus plant-ants. Biology Letters, 7(3), 475–479. doi:10.1098/rsbl.2010.0920

Russell, J. A., Moreau, C. S., Goldman-Huertas, B., Fujiwara, M., Lohman, D. J., & Pierce, N. E. (2009). Bacterial gut symbionts are tightly linked with the evolution of herbivory in ants. Proceedings of the National Academy of Sciences, 106(50), 21236–21241.

Russell, J. A., Sanders, J. G., & Moreau, C. S. (2017). Hotspots for symbiosis: Function, evolution, and specificity of ant-microbe associations from trunk to tips of the ant phylogeny (Hymenoptera: Formicidae). Myrmecological News, 24, 43–69.

Sanders, J. G., Łukasik, P., Frederickson, M. E., Russell, J. A., Koga, R., Knight, R., & Pierce, N. E. (2017). Dramatic differences in gut bacterial densities correlate with diet and habitat in rainforest ants. Integrative and Comparative Biology, 10.1093/icb/icx088. doi:10.1093/icb/icx088

Sanders, J. G., Powell, S., Kronauer, D. J. C., Vasconcelos, H. L., Frederickson, M. E., & Pierce, N. E. (2014). Stability and phylogenetic correlation in gut microbiota: lessons from ants and apes. Molecular Ecology, 23(6), 1268–1283. doi:10.1111/mec.12611

Sandström, J. P., & Moran, N. A. (2001). Amino acid budgets in three aphid species using the same host plant. Physiological Entomology, 26(3), 202–211.

Sandström, J., & Pettersson, J. (1994). Amino acid composition of phloem sap and the relation to intraspecific variation in pea aphid (Acyrthosiphon pisum) performance. Journal of Insect Physiology, 40(11), 947–955.

Seipke, R. F., Barke, J., Ruiz-Gonzalez, M. X., Orivel, J., Yu, D. W., & Hutchings, M. I. (2012). Fungus-growing Allomerus ants are associated with antibiotic-producing actinobacteria. Antonie van Leeuwenhoek, 101(2), 443–447. doi:10.1007/s10482-011-9621-y

Shin, S. C., Kim, S.-H., You, H., Kim, B., Kim, A. C., Lee, K.-A., … Lee, W.-J. (2011). Drosophila microbiome modulates host developmental and metabolic homeostasis via insulin signaling. Science, 334(6056), 670. doi:10.1126/science.1212782

Stamatakis, A. (2006). RAxML-VI-HPC: maximum likelihood-based phylogenetic analyses with thousands of taxa and mixed models. Bioinformatics, 22(21), 2688–2690. doi:10.1093/bioinformatics/btl446

Stoll, S., Gadau, J., Gross, R., & Feldhaar, H. (2007). Bacterial microbiota associated with ants of the genus Tetraponera. Biological Journal of the Linnean Society, 90(3), 399–412.

Tillberg, C. V., Holway, D. A., LeBrun, E. G., & Suarez, A. V. (2007). Trophic ecology of invasive Argentine ants in their native and introduced ranges. Proceedings of the National Academy of Sciences, 104(52), 20856–20861.

Van Treuren, W., Ponnusamy, L., Brinkerhoff, R. J., Gonzalez, A., Parobek, C. M., Juliano, J. J., … Meshnick, S. R. (2015). Variation in the microbiota of Ixodes ticks with regard to geography, species, and sex. Applied and Environmental Microbiology, 81(18), 6200–6209. doi:10.1128/AEM.01562-15

Ward, P. S., & Downie, D. A. (2005). The ant subfamily Pseudomyrmecinae (Hymenoptera: Formicidae): phylogeny and evolution of big-eyed arboreal ants. Systematic Entomology, 30(2), 310–335. doi:10.1111/j.1365-3113.2004.00281.x

Xu, H., Luo, X., Qian, J., Pang, X., Song, J., Qian, G., … Chen, S. (2012). FastUniq: A fast de novo duplicates removal tool for paired short reads. PLoS ONE, 7(12), e52249. doi:10.1371/journal.pone.0052249

Zientz, E., Feldhaar, H., Stoll, S., & Gross, R. (2005). Insights into the microbial world associated with ants. Archives of Microbiology, 184(4), 199–206. doi:10.1007/s00203-005-0041-0

